# Optimal confidence for unaware visuomotor deviations

**DOI:** 10.1101/2021.10.22.465492

**Authors:** Michael Pereira, Rafal Skiba, Yann Cojan, Patrik Vuilleumier, Indrit Bègue

## Abstract

Numerous studies have shown that humans can successfully correct deviations to ongoing movements without being aware of them, suggesting limited conscious monitoring of visuomotor performance. Here, we ask whether such limited monitoring impairs the capacity to judiciously place confidence ratings to reflect decision accuracy (metacognitive sensitivity). To this end, we recorded functional magnetic resonance imaging data while thirty-one participants reported visuomotor cursor deviations and rated their confidence retrospectively. We show that participants use a summary statistic of the unfolding visual feedback (the maximum cursor error) to detect deviations but that this information alone is insufficient to explain detection performance. The same summary statistics is used by participants to optimally adjust their confidence ratings, even for unaware deviations. At the neural level, activity in the ventral striatum tracked high confidence, whereas a broad network including the anterior prefrontal cortex encoded cursor error but not confidence, shedding new light on a role of the anterior prefrontal cortex for action monitoring rather than confidence. Together, our results challenge the notion of limited action monitoring and uncover a new mechanism by which humans optimally monitor their movements as they unfold, even when unaware of ongoing deviations.

## Introduction

Whether reaching for popcorn while viewing a movie or biking and enjoying the scenery, humans rely on reciprocal intricate connections between vision and motor processing to perform efficient behavior. Such visuomotor loops seem to occur mostly in the absence of awareness. Indeed, seminal work from (Fourneret & Jeannerod, 1998) showed that participants are unaware of their true hand position under imposed visuomotor deviations, although they appropriately correct all trajectories. Humans also neglect small spatial incongruences in feedback about their own movements (Farrer et al., 2008) and can reach targets that they cannot consciously report (Binsted et al., 2007) or that are displaced without them noticing (Goodale et al., 1986). These findings support the notion that participants show limited monitoring of the details of their movement, as long as their goal is achieved (Blakemore et al., 2002; Custers & Aarts, 2010; Gaveau et al., 2014)

A contradictory line of evidence comes from work on metacognition, i.e., the ability to monitor and control one’s internal processes (Flavell, 1979; Koriat, 2006). The standard measure of metacognition in humans is to ask them to make a decision and subsequently rate their confidence in the accuracy of that decision. Recent studies found that participants are able to adjust confidence ratings to their actual visuomotor decisions (Sinanaj et al., 2015; Locke et al., 2020; Charles et al., 2020; Arbuzova et al., 2020) indicating appropriate metacognitive monitoring of visuomotor performance. In contrast to Fourneret & Jeannerod ‘s results, these studies suggest that humans appropriately monitor their visuomotor performance.

Here, we set out to better understand this contradiction by comparing metacognitive sensitivity when participants explicitly report detecting deviations in their movements and when they do not. For this, we asked thirty-one participants to make straight reaching movements with a joystick, while lying in the MRI scanner. The speed of the cursor on the screen was set so that movements lasted two seconds. We introduced visual deviations to their trajectory in 79% of the trials by applying a clockwise or counter-clockwise visuomotor rotation of the joystick-to-cursor mapping (Fourneret & Jeannerod, 1998). Participants had to correct for these deviations, detect and report them (Farrer et al., 2008), and then rate their confidence in the accuracy of their detection responses on a scale ranging from 1 = not certain to 5 = completely certain (Figure 1A). We could thus determine how visual feedback of the participants’ movements is integrated into their confidence ratings, while dissociating trials for which they reported being aware of the deviations (hits) from those for which they were unaware of the deviation (misses). We then sought to identify brain regions whose activity correlated with confidence, as well as those implicated in action monitoring.

**Figure 1.**
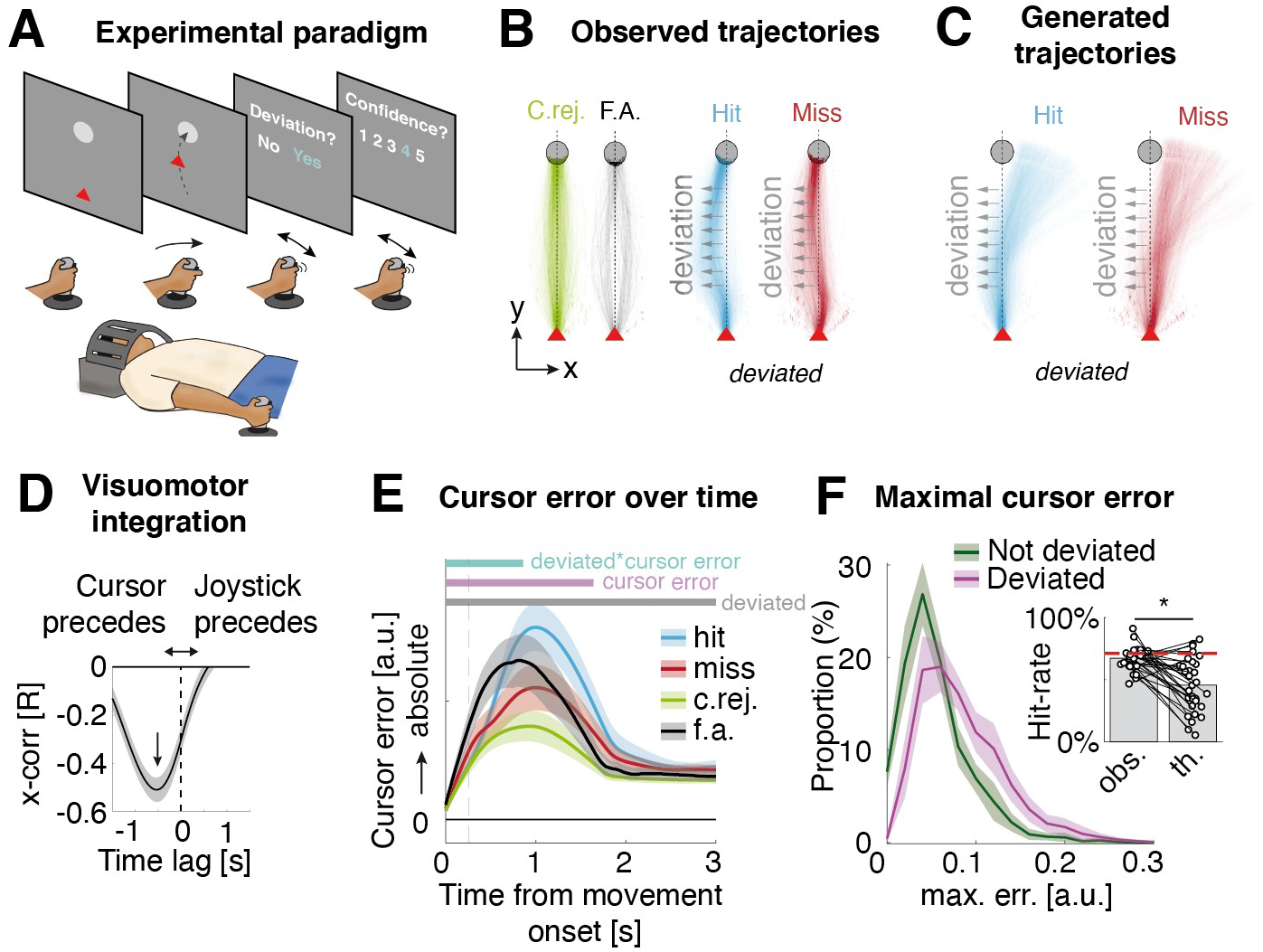
Experimental paradigm and detection responses. A) Participants started every trial by using the joystick to bring a visible cursor (red triangle) to a target (circle) on top of the screen. A deviation was applied to the cursor in 79% of the trials, titrated so as to reach 71% of detection accuracy. After reaching the target, participants reported whether the cursor was deviated or not (detection) and how confident they were about their answer. B) Trajectories of the cursor for correct rejections (green), false alarms (black), hits (blue) and misses (red). Note that deviations could be towards the left or the right, but right deviation trials are mirrored and pooled with left trials for display purposes. C) Generated trajectories obtained by recomputing the cursor position had there been no deviation for hits (blue) and misses (red). D) Cross-correlation between cursor error and joystick lateral position. The vertical arrow indicates the maximal cross-correlation (negative). E) Cursor error over time for hits (blue), misses (red), correct rejections (green) and false alarms (black). The significant (p < 0.05; FDR corrected) main effects of cursor error (respectively, of deviation) over time is depicted by the purple (respectively, grey) line; and the significant interaction effects between deviated trials and cursor error is depicted by the cyan line. F) Distributions of maximal cursor error for deviated (pink) and non-deviated (green) trials across participants (see Supplementary Figure 1 for distributions of individual participants). Inset: participants’ observed (obs.) hit-rates (HR) where compared to theoretical (th.) hit-rates derived from a receiving operating characteristic analysis of these distributions (Supplementary Figure 2). The horizontal dashed line represents the hit-rate target of the staircase procedure (71%). In all panels, shaded areas indicate 95% confidence intervals.

## Results

### Participants correct for deviations

For display purposes, we mirrored cursor trajectories for rightward deviations and combined them with leftward deviations, as there was no effect of the side of the deviation on detection responses (t(5164) = 1.33, p = 0.18; generalized linear mixed effect model with a binomial distribution). For both detected (hits) and undetected (misses) trials, all trajectories correctly ended on the target (Figure 1B) showing that participants always corrected for the experimentally induced deviations. Reconstructions of what trajectories would be without deviation showed clear corrective behavior for both hits and misses (Figure 1C). We also assessed the relation between joystick handle position (i.e. motor command) and cursor position (i.e. visual feedback). For this, we cross-correlated cursor and joystick position vectors across every trial, averaged over trials and participants and found that the strongest correlation occurred with a lag of −0.53 s ± 0.01 s (Figure 1D). This latency was similar between deviated (−0.54 s ± 0.02) and non-deviated trials (−0.49 s ± 0.07; t(30) = −0.66, p = 0.51), confirming that participants corrected cursor position through joystick adjustments, irrespectively of the presence of a deviation.

### Detection responses rely partly on maximal cursor error

A one-up/two-down staircase procedure titrated the angle of the deviation to an average of 19.87° ± 1.93, leading to 67.2% ± 1.6 correctly detected deviations (d’ = 1.60 ± 0.12). Participants were conservative in their response (c = 0.33 ± 0.07, t(30) = 4.86; p < 0.001) and made only few false alarms (7 ± 1 trials corresponding to 17.4% ± 2.8% of non-deviated trials). For deviated trials, small variations in the imposed deviation angle due to the staircase procedure had no effect on detection responses (t(5164) = 1.22, p = 0.22).

When analyzing data over time, we used the horizontal distance between the traced trajectories and the midline as an (unsigned) measure of cursor error (Figure 1E). We then regressed detection responses using cursor error and trial deviation (deviated vs. non-deviated) as fixed effects (generalized mixed effect models for every time point). We found an interaction effect during the first 0.89 s of the movement (p < 0.05; corrected for false-discovery rate (FDR) over time; Figure 1E), showing that during this time, the relation between detection responses and cursor error depended on whether the trials were deviated or not. Furthermore, there was a main effect of cursor error without an interaction in a time window ranging from 0.90 s to 1.64 s after movement onset (p < 0.05), suggesting that during this part of the movement, cursor error had an equal influence on detection responses, independently of whether the trial was deviated or not. There was an effect of deviated trials during the whole movement (p < 0.001), as this factor did not depend on time. It is worth noticing that cursor error peaked at inconsistent latencies across trials. We thus compared the best model (over time) with a single model using the *maximal* cursor error over time. The latter fitted the detection responses better (Deviance Information Criterion (DIC) relative decrease: 1.72 %; adjusted R^2^ = 0.79). The main effects for this model remained significant (maximal error: t(6403) = 5.31; p < 0.001; deviated trials: t(6403) = 7.49; p < 0.001) but there was no interaction (t(6403) = −1.32; p = 0.18), confirming that introducing deviations does not change the way visual feedback (quantified by maximal cursor error) is integrated into detection responses, but significantly biases detection (deviated trials are more correctly reported). Interestingly, the main effect of deviation suggests that participants possibly relied on some proxy to the deviations not contained in the visual feedback information alone.

To investigate this possibility further, we asked whether the maximal cursor error held enough information to explain detection performance. We attempted to decode deviated trials by setting a discrimination threshold on the maximal cursor error of individual participants (Figure 1F, Supplementary Figure 1), similarly to what participants would do if they were to rely only on visual feedback. We then selected the discrimination threshold that matched the observed false alarm rate and computed the theoretical hit-rate from the cursor error distribution (see Supplementary Figure 2 for the full receiving operating characteristic curve analysis). This theoretical hit-rate, corresponding to the theoretical performance that could be achieved using maximal cursor error alone was much lower (42 % ± 4) than the observed hit-rate (67% ± 2; t(30) = 5.16; p < 0.001; Figure 1F, inset), confirming that participants could not have reached the observed detection performance relying on visual feedback alone.

### Confidence also scales with cursor error and a proxy to deviation

We then turned to confidence ratings. Average confidence was higher for hits (4.11 ± 0.12) than for misses on deviated trials (3.69 ± 0.13; t(30) = 2.79; p = 0.0091), and even higher for correct rejections (4.63 ± 0.09) compared to hits (t(30) = 3.91; p < 0.001; Figure 2A). We regressed confidence ratings and compared different models including various summary statistics of cursor error over time (Figure 2B). We hereafter report the best model in terms of DIC, obtained by adding a predictor for maximal cursor error over time (max. err.) and for deviated trials. This model resulted in a 5.7 % relative decrease in DIC compared to a model including only the detection response regressor (Figure 2C; adjusted R^2^ = 0.29). The deviated trials x response interaction effect was significant (t(6399) = 6.13, p < 0.001), suggesting that participants still relied on a proxy to deviation not fully explained by maximal cursor error alone. This result is supported by the fact that models including trial deviation fitted the data better (Figure 2C). Moreover, there was an interaction between deviated trials and maximal cursor error (t(6399) = 2.29, p = 0.022) and between response and maximal cursor error (Figure 2D-E; t(6399) = 5.67, p < 0.001). There was no triple interaction between response, deviated trials and maximum cursor error (t(6399) = −1.81, p = 0.069). Alternative models, such as including the cursor error at the onset of the deviation (onset err.), the average (signed) cursor position (avg. pos.) or the average of the cursor error (avg. err.) yielded lower improvements in relative DIC. These results show that participants rated their confidence by conditioning the maximal cursor error to their detection response, with a confidence bias for not deviated trials that was dependent on their response (e.g. more confident for correct rejections vs misses; less confident for false alarms vs. hits).

**Figure 2.**
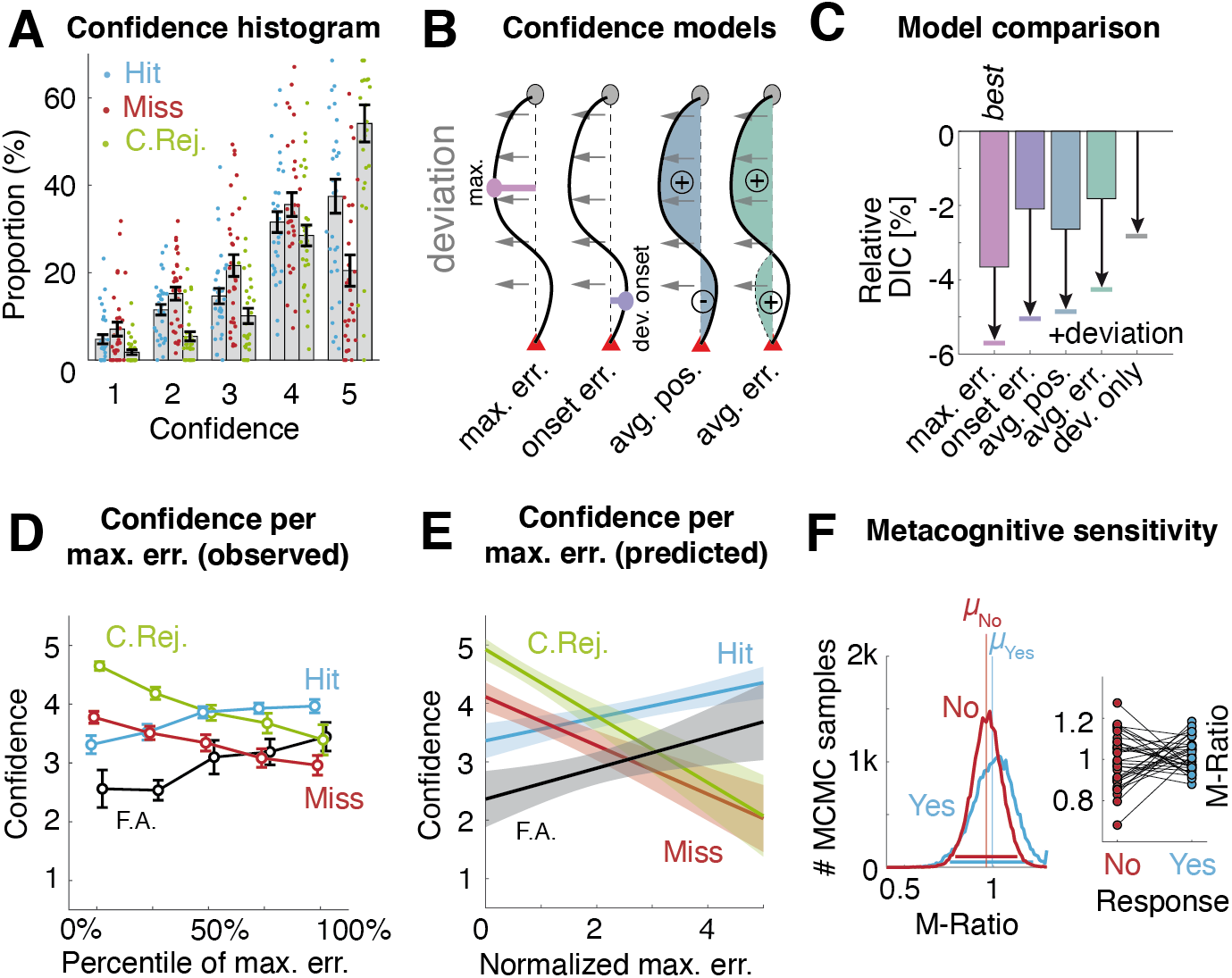
Confidence ratings and metacognition. A) Distribution of confidence ratings for hits (blue), misses (red) and correct rejections (green). Each point represents the data of one participant. B) Schematic depiction of the four regressors used for the four confidence models tested: max. err.: maximal cursor error; onset. err: cursor error at the onset of the deviation; avg. pos.: cursor position (signed) averaged along the trajectory; avg. err.: cursor error (unsigned) averaged along the trajectory. C) Relative improvement in DIC (compared to a model with no cursor information) for models without (bar plot) and with (horizontal line) a ‘deviated trial’ predictor. Note that the max. err. model shows the largest improvement. D) Confidence for different percentiles of maximal cursor error (max. err.) for hits (blue), misses (red) and correct rejections (green). E) Fixed effects predictions of confidence for comparable levels of max. err. (normalized per participants) for hits (blue), misses (red), correct rejections (green) and false-alarms (black). F) Hierarchical Bayesian estimation of response-specific metacognitive sensitivity using the M-Ratio. Left: posterior probability for yes (blue) and no (red) responses Vertical lines show the mean M-Ratio and horizontal bars show the 95% confidence interval. Right: Single participant estimates of the M-Ratio. In all panels, shaded areas and whiskers indicate 95% confidence intervals.

To better understand this relation between confidence and maximal cursor error, we fitted the data independently for deviated and non-deviated trials. We confirmed the interaction of response and maximal cursor error for both trial types (deviated: t(5162) = 6.30, p < 0.001; non-deviated: t(1237) = 5.24, p < 0.001). Importantly, and contrary to the hypothesis of limited monitoring, we also found that confidence was related to cursor error for all trial conditions (p < 0.01 for hits, misses, correct rejections, and false alarms). Together, these results show that confidence increases with maximal cursor error when participants report detecting the deviation but also decreases with maximal cursor error when participants do not report detecting the deviation, independently of whether the deviation was due to an experimental manipulation (e.g. during misses) or to their own intrinsic visuomotor variability (e.g. during correct rejections) (Figure 2D-E).

### Preserved metacognitive efficiency for unreported deviations

Finally, to confirm that participants correctly monitored their visuomotor actions, we measured their ability to use information available to the detection response in their confidence. For this, we fitted a response-specific hierarchical Bayesian model based on signal detection theory (Fleming, 2017; Maniscalco & Lau, 2014) which estimates metacognitive efficiency while controlling for task performance. This procedure estimates a meta-d’ measure, namely the d’ that would produce a similar distribution to that of the observed confidence ratings (Maniscalco & Lau, 2012). The output consists in the M-Ratio: a ratio between meta-d’ and d’, which quantifies metacognitive sensitivity, or how well information from first-order performance informs the metacognitive process. This analysis revealed a metacognitive efficiency of 0.96 for the “no” responses, indicating that participants used all information available to adjust their confidence ratings. Metacognitive sensitivity for “yes” responses was similar (0.99) confirming that reporting deviation does not improve metacognitive efficiency (Figure 2F).

### BOLD correlates of deviation detection and maximal cursor error

To investigate the neural substrate of the mechanisms described above, we used trial-by-trial measures (confidence, response, maximal cursor error and deviation angle) as parametric regressors to model the BOLD signal during each movement. All regressors showed only limited correlation (maximal absolute mean R = 0.41), similar to previous fMRI studies on confidence (Fleming et al., 2018). We found increased activity in the right primary visual cortex when participants detected the deviations (regressor for yes responses > 0; Figure 3B; Table 1), while the opposite contrast did not yield activity above statistical threshold. We found that larger maximal cursor error (parametric regressor for maximal cursor error > 0) yielded widespread BOLD activity increases in visuomotor and subthalamic regions, as well as in the left insula, right mid-cingulate and inferior frontal gyrii and lateral anterior prefrontal cortex (aPFC; Figure 1B; Table 1). Smaller cursor error yielded no activity beyond the statistical threshold.

**Figure 3.**
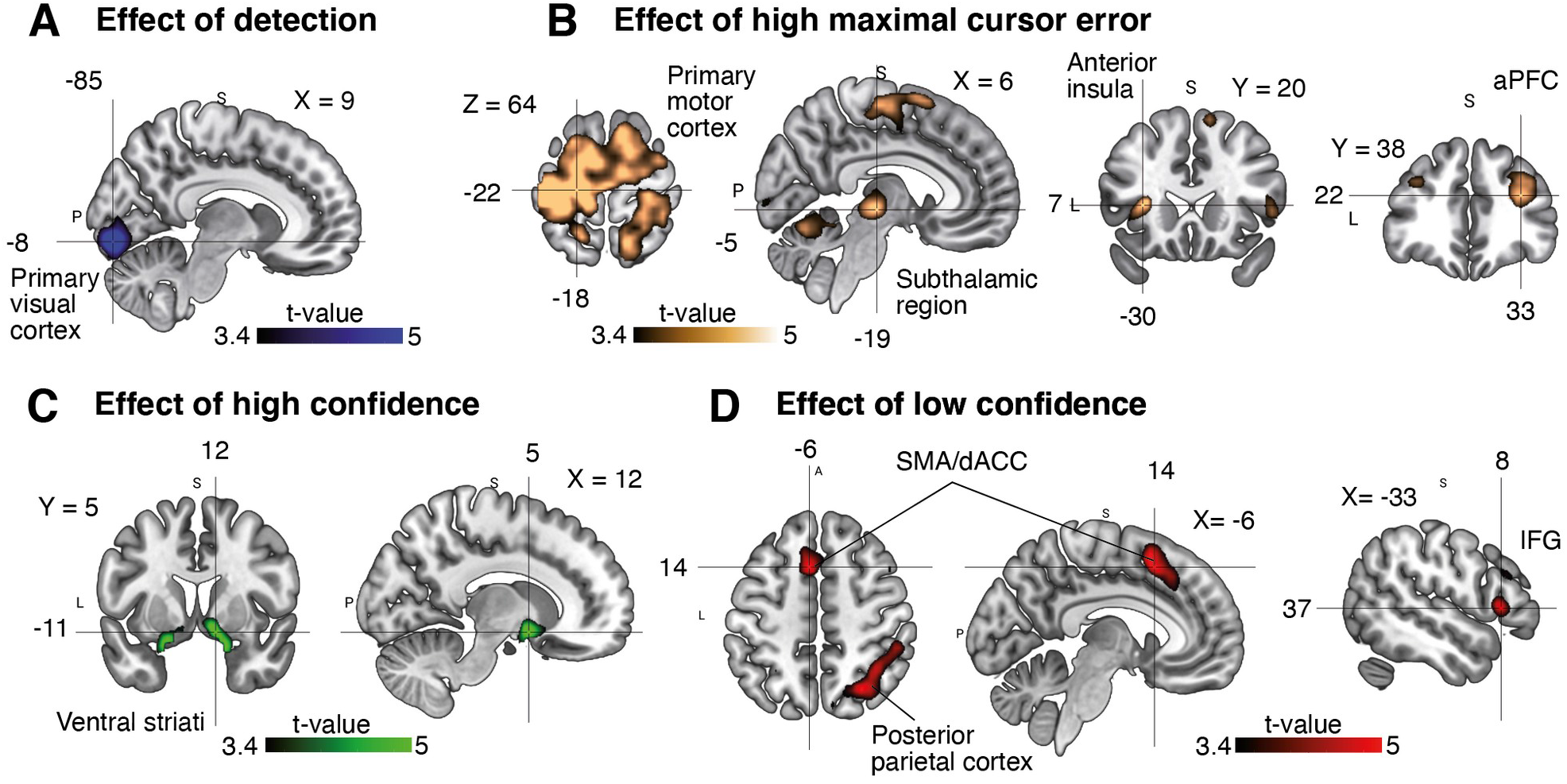
Statistical maps of parametric modulation contrast for A) explicit detection (yes responses), B) high maximal cursor error, C) high confidence and D) low confidence. Note that colors represent different parametric regressors and are independent from Figure 1. Results are displayed at p < 0.001 uncorrected. aPFC: anterior prefrontal cortex; SMA: supplementary motor area; dACC: dorsal anterior cingulate cortex. IFG: inferior frontal gyrus. See text Tables 1 and 2 for other brain activations.

**Table 1.** Summary of whole brain regions’ activation for response and maximal cursor error

|  |  |  |  |  |  |
| --- | --- | --- | --- | --- | --- |
| Brain region | MNI Coordinates<br>mm (xyz) |  |  | Cluster<br>extent<br>(voxels) | Significance |
| Yes responses |  |  |  |  |  |
| Right primary visual cortex | 9 | -85 | -8 | 602 | T= 6.64, $p_{FWEC} < 0.001$ |
| High maximal cursor error |  |  |  |  |  |
| Left primary motor cortex | -18 | -22 | 64 | 3563 | T= 8.18, $p_{FWEC} < 0.001$ |
| Right inferior occipital gyrus | 45 | -67 | 1 | 927 | T= 6.17, $p_{FWEC} < 0.001$ |
| Left anterior insula | -30 | 20 | 7 | 632 | T= 5.33, $p_{FWEC} < 0.001$ |
| Right subthalamic region | 6 | -19 | -5 | 285 | T= 5.26, $p_{FWEC} < 0.001$ |
| Right lateral anterior prefrontal cortex | 33 | 38 | 22 | 229 | T= 5.13, $p_{FWEC} < 0.001$ |
| Right mid-cingulate gyrus | 15 | -19 | 37 | 106 | T= 4.98, $p_{FWEC} = 0.028$ |
| Right inferior frontal gyrus | 48 | 29 | 1 | 279 | T= 4.47, $p_{FWEC} < 0.001$ |
| Left calcarine cortex | -12 | -85 | -8 | 147 | T= 4.41, $p_{FWEC} = 0.008$ |
| Left superior occipital gyrus | -21 | -79 | 28 | 108 | T= 4.16, $p_{FWEC} = 0.026$ |
| No surviving voxels for the contrasts: Low maximal error, No responses; $p_{FWEC} = p$ corrected for multiple comparisons at the cluster level | | | | | |

### BOLD correlates of confidence

We then turned to confidence and found that higher confidence (parametric regressor for confidence > 0) was related to increased bilateral ventral striatum activity, including the left amygdala (Figure 3C; Table 2). Lower confidence (parametric regressor for confidence < 0) was associated with increased activity in the left supplementary motor area (SMA), extending to the dorsal anterior cingulate cortex (dACC) and the left inferior and middle frontal gyri, as well as the right posterior parietal cortex (Figure 3D; Table 2).

**Table 2.**
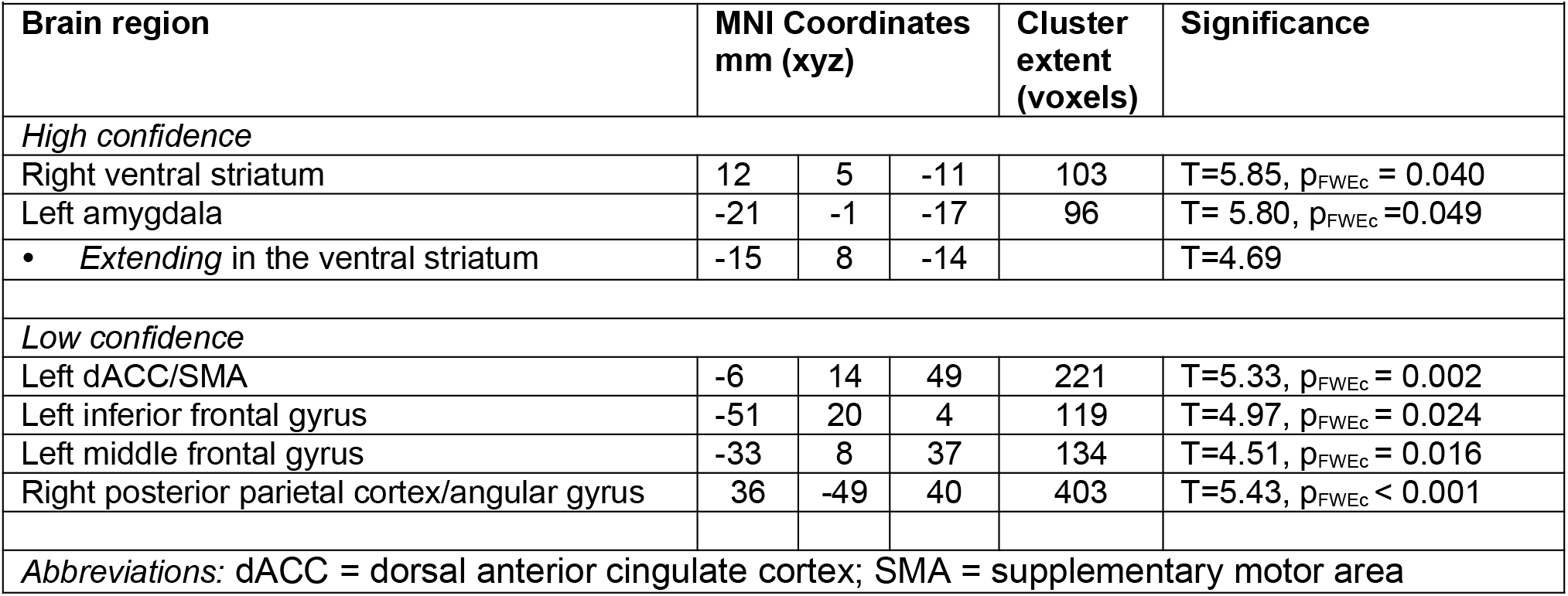
Summary of whole brain regions’ activation for confidence

| Table 2. Summary of whole brain regions' activation for confidence |  |  |  |  |  |
| --- | --- | --- | --- | --- | --- |
| Brain region | MNI Coordinates<br>mm (xyz) |  |  | Cluster<br>extent<br>(voxels) | Significance |
| High confidence |  |  |  |  |  |
| Right ventral striatum | 12 | 5 | -11 | 103 | T=5.85, p<sub>FWec</sub> = 0.040 |
| Left amygdala | -21 | -1 | -17 | 96 | T= 5.80, p<sub>FWec</sub> =0.049 |
| • Extending in the ventral striatum | -15 | 8 | -14 |  | T=4.69 |
| Low confidence |  |  |  |  |  |
| Left dACC/SMA | -6 | 14 | 49 | 221 | T=5.33, p<sub>FWec</sub> = 0.002 |
| Left inferior frontal gyrus | -51 | 20 | 4 | 119 | T=4.97, p<sub>FWec</sub> = 0.024 |
| Left middle frontal gyrus | -33 | 8 | 37 | 134 | T=4.51, p<sub>FWec</sub> = 0.016 |
| Right posterior parietal cortex/angular gyrus | 36 | -49 | 40 | 403 | T=5.43, p<sub>FWec</sub> < 0.001 |
| Abbreviations: dACC = dorsal anterior cingulate cortex; SMA = supplementary motor area |  |  |  |  |  |

## Discussion

We studied the behavioral and neural correlates of confidence for the detection of visuomotor deviations. We show that although participants did not report a third of the deviations introduced experimentally in the trajectory of their movement, metacognitive efficiency was not impaired by this lack of awareness; participants optimally calibrated their confidence judgments to the accuracy of their detection responses even when they reported no deviations. Furthermore, model selection revealed that whether participants were aware of the deviation or not, they relied on a summary statistic of visual feedback (i.e. maximal cursor error) for which we report the hemodynamic correlates. Importantly, participants also relied on a proxy to the deviations showing that observed detection performance could not be achieved based on visual feedback alone. Finally, we extend in the visuomotor domain previous findings of neural correlates of perceptual confidence (Hebart et al., 2016; Morales et al., 2018; Pereira et al., 2020; Rouault & Fleming, 2020; Vaccaro & Fleming, 2018), namely, neural localization of high confidence in the ventral striatum and of low confidence in the medial and lateral frontal cortex as well as in the posterior parietal cortex. However, we show that the aPFC, a region extensively linked with perceptual metacognition, modulates its activity as a function of visual feedback but not of confidence.

In our task, detection performance was kept fixed through an adaptive staircase procedure and the resulting small fluctuations in the angle of deviation had no significant effect on the detection responses, allowing us to examine confidence fluctuations independently from performance effects. We found a delay of 0.53 s between the joystick angle (*what the hand is doing)* and the cursor position (*what the eyes are seeing)*. These visuomotor correlations show that participants performed the task as required considering their joystick corrections where related to prior deviations in the trajectory. This delay was comparable between deviated and not deviated trials, showing that participants corrected externally – imposed deviations similarly to endogenous (i.e. internally-produced) errors in the non-deviated trajectories (e.g. Pereira et al., 2017).

In terms of detection responses, we found that participants report a deviation when their cursor error was high (hits and false-alarms) and to not report it when their cursor error was low (misses and correct rejections). However, we also found a significant effect of deviation *per se*. When comparing the amount of information contained in visual feedback and the detection performance, we found that participants could not possibly have relied on visual feedback alone. Other measures integrating cursor position over time yielded similar results. These findings suggest that to perform well at the task, participants must have at least partially relied on additional information, such as stemming from an internal model to compare their cursor position to a self-generated prediction, based on the efferent copy of their motor command (Kawato, 1999; Wolpert et al., 1995). The later assumption agrees well with past research on the sense of agency (Haggard, 2017). We therefore hypothesize that participants applied a strategy involving a weighted average of both the feedback prediction error and visual feedback monitoring for efficient task-based performance.

To further assess the extent to which participants monitor their actions, we examined participants’ metacognitive sensitivity when they reported being unaware of the deviations that they successfully corrected. Surprisingly, metacognitive sensitivity was close to 1 for both aware and unaware deviations, suggesting that the evidence available for confidence was similar to that available for the detection report, i.e. no additional metacognitive noise (Maniscalco & Lau, 2012; Shekhar & Rahnev, 2021). Even though participants did not report some deviations, their confidence ratings still discriminated deviated from non-deviated trials in an optimal manner (considering the information available for detection). Moreover, when disentangling the factors influencing confidence, we found that confidence increased with maximal cursor error when participants reported the deviation and, crucially, decreased with maximal cursor error when participants did not report the deviation. This behavior reveals a judicious use of confidence, irrespective of the awareness of the deviation. Furthermore, participants also appeared to integrate in their confidence ratings additional information such as feedback prediction errors. We thus argue that participants have good monitoring of their motor actions, having access to at least a summary of their motor behavior for conservative detection and optimal confidence. Therefore, our findings question the notion of limited monitoring put forth by (Fourneret & Jeannerod, 1998) and others (Blakemore et al., 2002; Desmurget & Sirigu, 2009): participants are unaware of some deviations that they correct, but they still have a ‘feel’ for their performance. A difference between Fourneret & Jeannerod’s study and ours is that instead of asking participants to report on the true position of their hand, we asked them to report on the presence or absence of a deviation (as in Farrer et al., 2008). One speculative explanation is that participants are only able to monitor the magnitude of their feedback prediction errors, which would explain why they can optimally rate their confidence but not report the actual position of their hand.

Our result of preserved metacognition for “no” responses also sharply contrast with studies on visual perception which describe lower metacognition for unaware stimuli (Kanai et al., 2010; Mazor et al.,2020; Pereira et al., 2021). Considering that the nature of metacognitive inefficiency is still unknown (Shekhar & Rahnev, 2021), we can only speculate on why metacognitive performances does not decrease for unaware deviations in our study. Confidence for unaware stimuli was proposed to depend on monitoring attention instead of perceptual evidence (Kanai et al., 2010; Mazor et al., 2020). According to this view, confidence for aware and unaware stimuli is based on different mechanisms. In other studies, reduced metacognitive efficiency for unaware stimuli was modeled using a single mechanism (Kellij et al., 2021; Pereira et al., 2021), based on the fact that the variance of the noise and stimuli (signal) differ. It could thus also be possible that in our study, smaller differences in variances between the noise and the signal allow metacognition to be preserved for “no” responses.

At the neural level, our fMRI results showed extended BOLD activations for increasing maximal cursor error in sensorimotor regions (especially contralateral to the moving hand; all participants were right-handed), occipital, anterior prefrontal and insular cortices, subthalamic regions as well as in the mid-cingulate and inferior frontal gyrus. These results suggest the existence of widespread action-monitoring processes (Limanowski et al., 2017). The aPFC has been extensively linked to perceptual metacognition using voxel-based morphometry (Fleming et al., 2010), TMS (Rahnev et al., 2016; Shekhar & Rahnev, 2018) or lesion studies (Fleming et al., 2014). Surprisingly, although BOLD activity in this region was shown to relate negatively with confidence in previous fMRI studies (Fleming et al., 2012; Fleming et al., 2018; Mazor et al., 2020; Pereira et al., 2020), in our study, the variance of the BOLD signal in the aPFC was explained by the maximal cursor error rather than by the confidence regressor. This result implies that the aPFC might not be involved in confidence *per se (how confident am I in the accuracy of my decision)* but rather in monitoring the performance of our actions (*how well am I performing this reaching movement)*. This view is consistent with a recent fMRI study where activity in the aPFC was only related to confidence when participants enacted their decisions with a motor action but not when they covertly rated their confidence in someone else’s decisions (Pereira et al., 2020). In line with our interpretation, patients with prefrontal lesions reported fewer deviations than healthy control despite similar corrective behavior using a similar task to ours (Slachevsky et al., 2001). In sum, our finding of increased aPFC activity during increased cursor error and not confidence thus pleads in favor of a novel role for the aPFC as a key region for monitoring action performance rather than the accuracy of decisions.

We found that low confidence related to activity in the medial frontal cortex, the left inferior and middle frontal gyri, and the right posterior parietal cortex, providing novel support in the visuomotor domain for fronto-parietal regions’ role in metacognitive processes via graded confidence computation (Hebart et al., 2016; Morales et al., 2018; Pereira et al., 2020; Rouault & Fleming, 2020; Vaccaro & Fleming, 2018). High confidence in visuomotor decisions engaged the ventral striatum, also corroborating fMRI findings in the perceptual domain (Hebart et al., 2016; Guggenmos et al., 2016; Rouault et al., 2018; Mazor et al., 2020) or clinical obsessive-compulsive disorder cohorts undergoing deep brain stimulation (de Haan et al., 2015; further discussed in Kiverstein et al., 2019). Apart from its well-known involvement in reward-based learning (Daniel & Pollmann, 2014), the ventral striatum also computes pseudo-reward prediction errors – defined as reward predictions errors related to the subjectively perceived progress in a given task – rather than merely to external (e.g. monetary) reward (Westbrook et al., 2016). It has been shown that these pseudo-reward prediction errors bias choice behavior, even in the absence of monetary reward (Mas-Herrero et al., 2019). Our results can therefore easily be reconciled with a putative role of the ventral striatum for valuation information predicting reward (Schultz et al., 1992; Pagnoni et al., 2002; Daniel & Pollmann, 2014), whereby in the absence of feedback, the valuation information corresponds to confidence (Daniel & Pollmann, 2012). Taken together, our findings support a key role of the ventral striatum in monitoring decisional signals for confidence (Daniel & Pollmann, 2012; Hebart et al., 2016; Vaccaro & Fleming, 2018) that can be used to adapt subsequent behavior in absence of external feedback (Guggenmos et al., 2016). In keeping with this, confidence in a perceptual task in a large non-clinical sample was negatively correlated to apathy, a psychopathological manifestation of a reduction in goal-directed behavior (Roualt et al. 2018). Future research initiatives extending metacognition research in clinical populations manifesting with deficits of goal-directed behaviors (e.g. negative symptoms of schizophrenia) should prove useful to better dissect confidence contribution in the underlying pathophysiology.

To conclude, we uncovered a plausible mechanism for the monitoring of visuomotor deviations: participants base their detection and confidence reports on the monitoring of a summary statistic of the cursor position but also by possibly comparing visual feedback to self-generated predictions. We mapped this monitoring and correcting of the cursor position to an extended network of brain regions including the aPFC, shedding light on a different role for this region than simply tracking confidence, that is monitoring action performance. Importantly, although participants did report being unaware of some deviations, their confidence ratings were as informative of their performance as when they reported deviations and monitoring the same summary statistics. Our results offer a plausible explanation for a paradox: that humans perform corrective actions in the absence of awareness but are good at attributing actions to themselves or to an external agent (Vignemont & Fourneret, 2004). Instead, we argue that even if participants are unaware of their corrections, they still can monitor their performance through some summary statistic. They only become aware that something is wrong when that summary statistic exceeds what could be expected from their own intrinsic motor variability. This has important implications as deficits in the awareness of action have been extensively linked to psychiatric diseases (Blakemore & Frith, 2003) such as schizophrenia (Frith et al., 2000; Voss et al., 2010). Our methodology should catalyze future research efforts in the visuomotor domain assessing whether schizophrenia, or more generally, psychosis spectrum patients have a metacognitive deficit (Rouy et al., 2021). It will be important to examine whether these clinical populations employ the same mechanisms to compute confidence as we describe here, and if so, how deficits in such mechanisms can be mapped onto specific pathophysiologic dimensions (positive i.e. psychotic and negative i.e. amotivational symptoms).

## Acknowledgements and funding

MP is supported by two Postdoc.Mobility fellowships from the Swiss National Science Foundation (P2ELP3_187974; P400PM_199251). IB is supported by a PRD fund from the University Hospital of Geneva (PRD 22-2020-1) as well as a Scientific chief resident fellowship from the University of Geneva, Switzerland. This study was conducted on the imaging platform at the Brain and Behavior Lab (BBL), at the University of Geneva, Switzerland and benefited from support of the BBL technical staff. Special thanks to Christophe Mermoud for programming of the experimental protocol. We thank Nathan Faivre for constructive comments on the manuscript.

## Methods

### Participants

We recruited thirty-two healthy right-handed participants. One participant did not complete the experimental task, therefore, the final sample included 31 participants (age: 26 years ± 4.7). Participants gave written informed consent prior to the experiment and received 20 Swiss francs per hour as compensation. They had normal or corrected-to-normal vision and reported no neurological or psychiatric disorder. The study was approved by the Ethics Committee of the University of Geneva and University Hospitals of Geneva (CER:11-214/NAC 11-077). All participants read and signed an informed consent form, and were screened for contraindications to MRI with a standard safety questionnaire. Structural analysis from this cohort has been already published (Sinanaj et al., 2015). Ten participants have been used as matched controls for a study on conversion disorders (Bègue et al., 2018).

### Experimental procedure

We asked participants to perform a visuomotor conflict-inducing task (Bègue et al., 2018; Sinanaj et al., 2015) adapted from a classic paradigm from (Fourneret & Jeannerod, 1998; Farrer et al., 2008). After a short preparation period (white triangle became red, duration 1-2 seconds, jittered) participants had to push the joystick handle forward in order to start moving towards a centrally-located target in the upper section of the screen. After reaching the target, participants reported whether they noticed any externally originating deviations of their trajectory (detection report), and subsequently rated their confidence in their own judgment on a scale ranging from 1 = not certain to 5 = completely certain. Participants did not receive feedback about the accuracy of neither detection, nor confidence judgments. We encouraged participants to use the whole confidence scale. The experimental manipulation consisted in introducing deviations of the visual trajectory on the screen, towards either the right or left side. These deviations were gradually applied (0.3 s ramp), starting after participants reached a fixed distance from the starting point corresponding to 13% of the vertical distance between the initial cursor error and the final target. Participants were informed that these externally originating deviations would not occur all the time, however, when they occurred, participants had to correct for these deviations in order to reach the target. Right/leftward corrections were possible through right/left pushes of the joystick handle on the left or right, respectively. Participants selected “Yes” or “No” responses through joystick handle movements on the right and left, respectively, then pressing a button.

We asked participants to perform a training run outside the scanner, consisting of 30 non-deviated trajectories to familiarize them with the joystick and experimental environment. For each experimental session, we ran an adaptive staircase procedure (Levitt, 1971) that made the task more difficult after two consecutive correct responses by increasing the next deviation by 2.64°, but made it easier after an incorrect response by reducing the next deviation by 1°. After the training session, participants entered the scanner and performed a ‘threshold’ session of 80 trials in order to stabilize the staircase procedure (data not analyzed). After the threshold session, participants completed two experimental runs. A structural T1 image was acquired between these two runs. Overall, there were 208 trials (21% without trajectory deviation). Each trial lasted 11.5 s and was followed by a blank screen with a jittered duration (3 to 6 s).

### Behavioral analyses

We excluded the first 80 trials until the adaptive staircase procedure converged. We defined a trial with a deviation that was reported as such by participants as a *hit* and as a *miss* in case it was not reported. A trial was a *correct rejection* when there was no deviation and participants correctly reported no deviation and a *false alarm* if participants reported a deviation. We grouped hits and false alarms into *“Yes”* responses and misses and correct rejections into *“No”* responses. We computed the sensitivity d’ and criterion c using signal detection theory (Stanislaw & Todorov, 1999). *Cursor error* was defined as the horizontal distance between the cursor position and the midline between the starting point (triangle at a lower central position on the screen) and the target (top central position). We then defined the *maximal cursor error* (max. err.) as the maximum of the cursor error the course of a trial. To avoid selecting the last sample (that might not lead to a correction, we preferred the highest peak of the cursor error rather than the maximum. These only differed in an average of 5.84 ± 0.89 trials per participants and this choice did not affect the results. *Onset error* (onset err.) was defined as the cursor error at the onset of the deviation. We also defined the *average position* (avg. pos.) as the absolute value of the average position of the cursor with respect to the sagittal line (the later can be negative) as well as the *average cursor error* (avg. err.) as the average of the distance between the cursor and midline (always positive). A trial with a large deviation to the right followed by a large deviation to the left would thus have an average position close to zero but a high average cursor error.

To build two-dimensional histograms of the cursor trajectories (Figure 1B, C), we mirrored trajectories when the deviation was towards the right (only for hits and misses). We then computed two-dimensional histograms of the position of the cursor in all trials of a condition (hit, miss, correct rejection and false alarms) and normalized the resulting histograms by the number of trials in that condition. Finally, we averaged across participants. For statistics, we defined (generalized) linear mixed effect models to analyze detection responses and confidence ratings. To regress detection responses, we used a binomial distribution with a logistic link function. Inclusion of random effects was guided by model selection based on deviance information criterion and led to the inclusion of all factors and interactions as random effects. All statistical tests were two-tailed. To assess the amount of information in the visual feedback, we performed receiving operator characteristics analyses by sliding a criterion along the maximum cursor error while computing the true- and false-positive rate which we plotted in Supplementary Figure 2. We then searched for the criterion leading to the same false-positive (false-alarm) rate as found in the data and compared the corresponding true-positive rate (hit rate) to the one observed in the data.

For Figure 1D, we binned the maximal cursor error into five quantiles computed independently for each participant but for all conditions together. For Figure 1E, we normalized maximal cursor error by its standard deviation computed over all conditions. To estimate metacognitive sensitivity, we used the ratio between meta-d’ (Maniscalco & Lau, 2014) and d’ estimated using the response-specific version of the HMeta-d’ toolbox (Fleming, 2017). We used the default parameters of three chains of 10’000 samples with 1’000 *burn in* samples for the MCMC procedures with no thinning. Visual inspection of MCMC and R_hat values well under 1.1 indicated good convergence.

### fMRI data collection, preprocessing and analyses

We acquired functional MRI images with a 3T whole-body scanner (Trio TIM, Siemens, Germany) with a 12-channel head-coil. Functional images were acquired with a susceptibility weighted EPI sequence with the following parameters: TR/TE = 2100/30 ms, flip angle = 80 degrees, PAT factor = 2, 64 × 64 voxel, 3.2 × 3.2 mm, 36 slices, 3.2 mm slice thickness, 20% slice gap. We acquired structural images using a T1-weighted 3D sequence using the following parameters: MPRAGE, TR/TI/TE = 1900/900/2.32 ms, flip angle = 9°, voxel dimensions: 0.9 mm isotropic, 256 × 256 × 192 voxels. We presented task stimuli on a back-projection screen inside the scanner bore using an LCD projector (CP-SX1350, Hitachi, Japan). We recorded responses via buttons placed on the joystick used for the visuomotor reaching task (HH-JOY-4, Current Designs Inc., USA

We used the SPM8 software (http://www.fil.ion.ucl.ac.uk/spm/) for statistical analyses of functional data with a standard pipeline. We first corrected for head movements between scans by an affine registration (Friston et al., 1995) and realignment to the mean of all images. The anatomical image was spatially normalized on the EPI template. The functional images were also normalized to the EPI template, which were thereby transformed into standard stereotaxic space and resampled with a 3 × 3 × 3 mm voxel size. The normalized images were spatially smoothed using an 8 mm full-width at half-maximum (FWHM) Gaussian kernel. We used the general linear model (GLM) framework implemented in SPM to analyze our data. We modeled the convolved standard hemodynamic response function with a delta (or “stick”) function at the onset of the preparatory phase (appearance of white triangle – PREP regressor), at the onset of the joystick movement (start of movement – MOV regressor), and at the onset of the response screen (yes vs. no, RESP regressor). To examine brain regions whose activity fluctuated with trial-by-trial confidence, we took a parametric modulation approach (Pereira et al., 2020): the “MOV” regressor event regressors were modulated by additional parametric factors representing the trial-by-trial values of confidence (CONF parametric modulator), maximal cursor error (ERR parametric modulator), response yes vs. no (YN parametric modulator) and angle of deviation (ANG). To account for head motion-related variance, we included the six differential parameters derived from the realignment process [x, y, and z translations (in millimeters) plus pitch, roll, and yaw rotations] as regressors of no interest. Low-frequency signal drifts were filtered using a cut-off period of 128 s. Global scaling was applied, with each fMRI value rescaled to a percentage value of the average whole-brain signal for that scan.

Contrast images from one-sample t-tests corresponding to each event (PREP, MOV, RESP) and their parametric modulators (CONF, ERR, YN, ANG), were fed into a second-level random-effect analysis. All second-level results are reported at a significance-level of p < 0.05 using cluster-extent family-wise error (FWE) correction with a voxel-height threshold of p < 0.001. In Figure 3, activations are displayed at a cluster-size threshold of 30 voxels, using MRIcroGL (http://www.cabiatl.com/mricrogl/). Data and analysis scripts from this study will be made freely available upon acceptance.

## Notes

**Conflict of interest** The authors declare no competing interests.

### Competing Interest Statement

The authors have declared no competing interest.

