## Supplementary materials for "Optimal confidence for unaware visuomotor deviations"

**Affiliations**
1. Université Grenoble Alpes, CNRS, LPNC UMR 5105, Grenoble, France
2. Laboratory for neurology and imaging of cognition, Department of Basic Neuroscience, University of Geneva, Geneva, Switzerland
3. Adult psychiatry division, Department of psychiatry, University hospitals of Geneva, Geneva, Switzerland
4. Laboratory for clinical and experimental psychopathology, Department of psychiatry, University of Geneva, Geneva, Switzerland

**Corresponding author:**
Indrit Bègue <>
Laboratory for clinical and experimental psychopathology, Department of psychiatry, University hospitals of Geneva, University of Geneva, Geneva, Switzerland

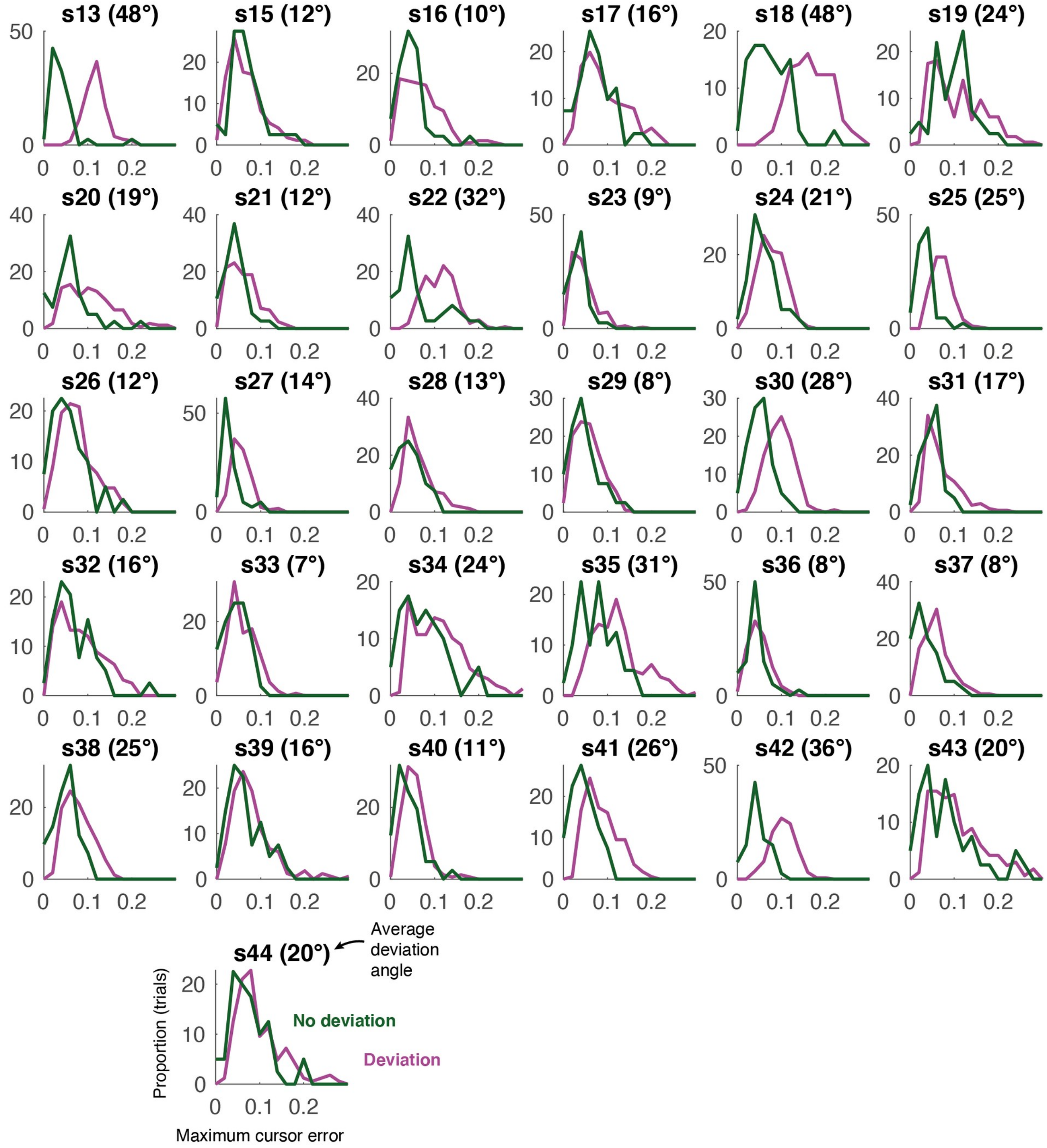


***Supplementary Figure 1: Distribution of maximal cursor error.***

*For deviated (light blue) and non-deviated (dashed black) trials for every participant. The average deviation angle (i.e. easiness of the task) obtained through the staircase procedure is displayed in the title of each panel. Only participants who failed to reach a low average deviation angle showed highly separable distributions (e.g. s13, s18, s42).*

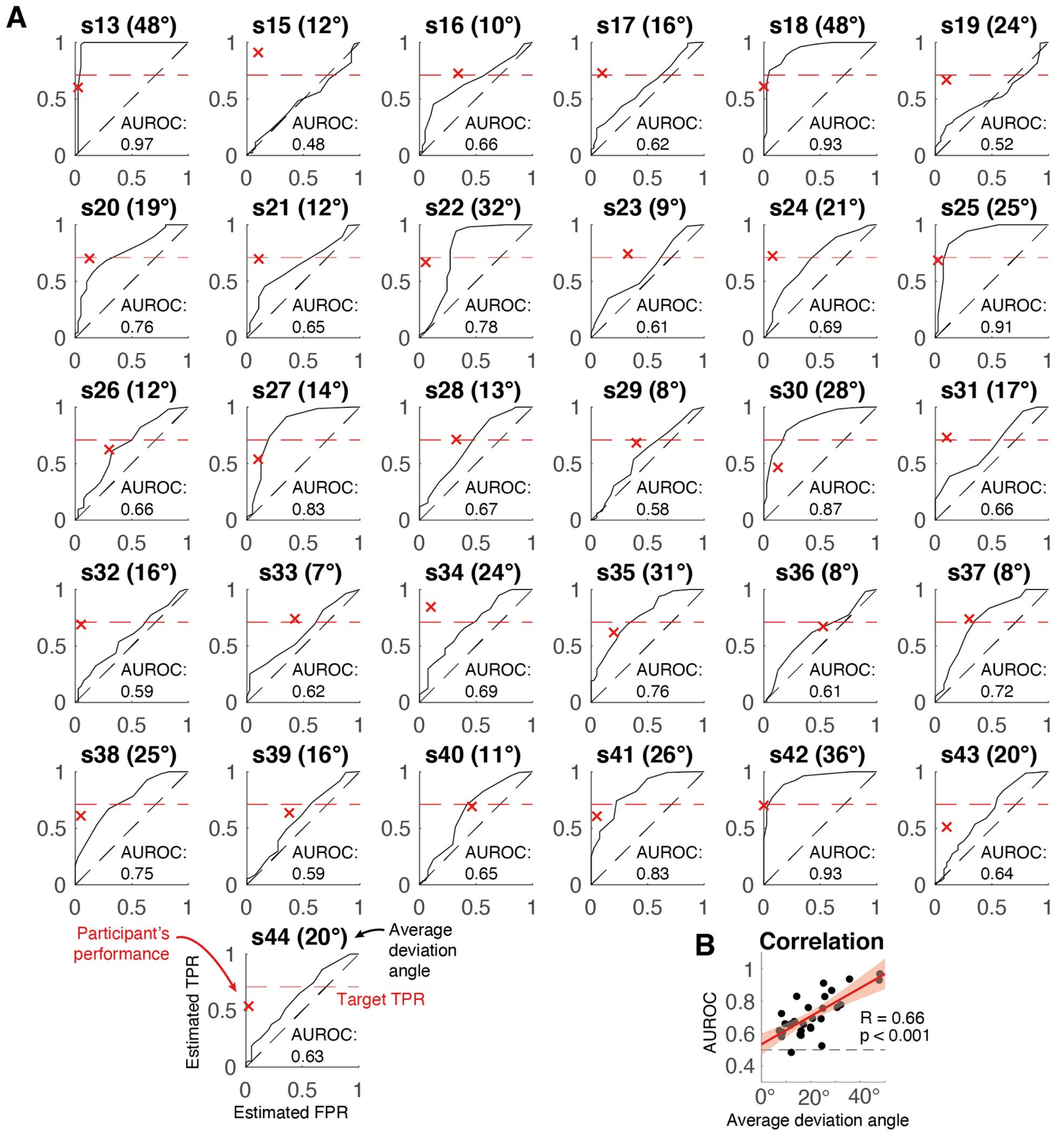


***Supplementary Figure 2: Receiving operating curve (ROC) analysis.***
*A) To compare individual participants' performances against the information contained in the maximal cursor error, we estimated the ROC curve (black trace) by sliding the criterion along the horizontal axis in Supplementary Figure 1 while computing true positive rate (estimated TPR) and false positive rate (estimated FPR). The dashed black trace represents the theoretical ROC curve for random performance (i.e. for non-separable distributions of maximal cursor error between deviated and non-deviated trials). We then displayed the actual performance from each participant in terms of TPR and FPR (red cross). If participants would rely only on the maximal cursor error to detect deviations, we would expect their performance to be under (or close to) the estimated ROC curve. This was the case only for a few participants (e.g. s30, s36, s40). These results indicate that most participants used additional information to detect deviations. We obtained very similar results using other indices such as cursor error at deviation onset, averaged cursor position, and averaged cursor error, suggesting that participants did not use another more integrative strategy either, even based on other visual cues from the cursor. B) Between-participant correlation between the estimated area under the ROC curve (AUROC) quantifying the information contained in the maximal cursor error and the average deviation angle achieved by the staircase procedure. This relation shows that the less good participants were at the task (high deviation angle), the more information they could use from the maximal cursor error.*
